# *OsPATROL1* overexpression accelerates stomatal opening to enhance photosynthetic induction and growth under fluctuating light in rice

**DOI:** 10.64898/2026.09.01.748733

**Authors:** Naoya Katsuhama, Ryutaro Morita, Yu Wakabayashi, Sangil Park, Hiroshi Fukayama, Naohiro Aoki, Ichiro Terashima, Wataru Yamori

## Abstract

Slow stomatal opening after increases in irradiance constrains carbon gain under fluctuating light, yet stomatal kinetics remain an underexplored target for crop improvement. Here, we investigated *Oryza sativa PROTON ATPASE TRANSLOCATION CONTROL 1* (*OsPATROL1)*, which encodes a Munc13-like protein implicated in stomatal regulation in *Arabidopsis thaliana*. *OsPATROL1* overexpression had modest, condition-dependent effects on steady-state gas exchange and did not alter stomatal morphology or biochemical traits. In contrast, it consistently accelerated stomatal opening and photosynthetic induction, reducing the stomatal conductance time constant during induction by 41–43%. During 12 h of simulated natural fluctuating light, *OsPATROL1*-overexpressing plants maintained higher stomatal conductance and net CO₂ assimilation rate, increasing cumulative assimilation by 8–12% while maintaining their intrinsic water-use efficiency (*iWUE).* Under artificial fluctuating light, overexpression alleviated growth reductions relative to steady light. Under glasshouse conditions, total biomass increased by 34–44%, accompanied by greater tiller number, root biomass, bleeding sap rate, and leaf nitrogen content. Taken together, these results indicate that *OsPATROL1* overexpression accelerates stomatal opening, enhances photosynthetic induction and daytime carbon gain without compromising *iWUE*, and is associated with greater growth.

## Introduction

Leaves within field canopies experience rapid fluctuations in irradiance caused by changes in the solar angle, cloud movement, wind, and mutual shading, resulting in strong spatial and temporal variation in photosynthetic photon flux density (PPFD) at the leaf surface (Pearcy, 1990; Nishimura et al., 1998; Yamori, 2016). Following an increase in irradiance, the net CO₂ assimilation rate (*A*) often remains below its steady-state value because stomatal opening and photosynthetic metabolism activation lag behind changes in light. Both stomatal and biochemical processes contribute to photosynthetic induction, and their relative importance changes over the course of induction (Kirschbaum and Pearcy, 1988; Pearcy, 1990). Such delays reduce the daily carbon gain and may ultimately constrain crop productivity under dynamic light environments (Long et al., 2015; Tanaka et al., 2019; Lawson and Matthews, 2020). Therefore, rice (*O. sativa*), a staple crop for more than half of the world’s population, is an important target for improving photosynthetic performance in dynamic light environments (Fukagawa and Ziska, 2019). Rice genotypes exhibit substantial natural variation in photosynthetic induction, indicating considerable scope for improvement (Acevedo-Siaca et al., 2020; Taniyoshi et al., 2020; Xiong et al., 2022; Taniyoshi et al., 2025). For example, the high-yielding cultivar Takanari exhibits faster increases in stomatal conductance (*g*_sw_), electron transport, and *A* than Koshihikari and achieves greater daily CO₂ assimilation under simulated natural fluctuating light (Adachi et al., 2019). More broadly, faster stomatal opening is associated with faster photosynthetic induction among diverse rice genotypes, although the relative contributions of stomatal and non-stomatal limitations vary among genotypes and induction phases (Acevedo-Siaca et al., 2020; Xiong et al., 2022; Taniyoshi et al., 2025). Taken together, these findings indicate that photosynthetic induction is an exploitable trait for increasing carbon gain under field-relevant irradiance fluctuations.

For the genetic improvement of dynamic photosynthesis in crops, photochemical and biochemical processes have been primarily targeted. Manipulation of the Rieske FeS protein, RuBP-regeneration enzymes, Rubisco content, or Rubisco activase activity has accelerated photosynthetic induction and increased productivity of tobacco, sorghum, sugarcane, or rice (López-Calcagno et al., 2020; Ermakova et al., 2023; Salesse-Smith et al., 2025; Yamori et al., 2026). These studies demonstrate the potential of manipulating non-stomatal processes; however, stomatal limitation remains an important target after prolonged exposure to low irradiance or during the later phases of induction (Kirschbaum and Pearcy, 1988; Pearcy and Seemann, 1990; Yamori et al., 2020; Taniyoshi et al., 2025).

Manipulating stomatal responses offers a complementary strategy, but increasing *g*_sw_ to promote CO₂ diffusion may also increase transpirational water loss. In rice, loss of *SLOW ANION CHANNEL-ASSOCIATED 1* (*SLAC1*), a guard-cell anion channel required for stomatal closure, causes constitutively high *g*_sw_ (Kusumi et al., 2012). This elevated *g*_sw_ accelerates photosynthetic induction and increases daily carbon gain under fluctuating light; however, it does not increase shoot biomass under fluctuating light and may impose a cost through excessive water loss (Yamori et al., 2020). Overexpression of *OSA1*, which encodes a plasma membrane H⁺-ATPase, enhanced steady-state *g*_sw_ and *A* and increased grain yield, but instantaneous water-use efficiency was reduced by 13–21% (Zhang et al., 2021). Conversely, overexpression of *OsNHX1* accelerated stomatal closure following reductions in irradiance and increased biomass and grain yield under drought, demonstrating the potential of manipulating closing kinetics to conserve water (Qu et al., 2020). In *Arabidopsis thaliana*, accelerating both stomatal opening and closure using the optogenetic K⁺ channel BLINK1 increased biomass under fluctuating light without increasing whole-plant water use (Papanatsiou et al., 2019). Collectively, these studies not only demonstrate the potential of manipulating stomatal behavior but also highlight the importance of matching *g*_sw_ to the changing photosynthetic demand rather than simply maintaining greater stomatal opening.

Plasma membrane H⁺-ATPases play a central role in light-induced stomatal opening. Blue light activates guard-cell H⁺-ATPases through phosphorylation of their C-terminal regulatory region, initiating the membrane hyperpolarization and ion uptake required for stomatal opening (Kinoshita and Shimazaki, 1999). Rather than directly increasing H⁺-ATPase expression, PROTON ATPASE TRANSLOCATION CONTROL 1 (PATROL1), a Munc13-like protein in *A. thaliana*, coordinates complementary guard- and subsidiary-cell responses through the stimulus-dependent trafficking of the plasma membrane H⁺-ATPase AHA1, thereby contributing to stomatal opening in response to light and CO₂ (Hashimoto-Sugimoto et al., 2013; Higaki et al., 2014). *PATROL1* overexpression accelerates stomatal opening and photosynthetic induction under fluctuating light, increasing photosynthesis and biomass while maintaining water-use efficiency (Kimura et al., 2020). Such stimulus-responsive control of H⁺-ATPase localization could help rapidly increase *g*_sw_ when CO₂ demand rises without constitutively maintaining high *g*_sw_.

Despite these findings in *A. thaliana*, whether PATROL1 has conserved physiological functions in crop species remains unknown. Rice and *A. thaliana* possess morphologically distinct stomatal complexes: rice has dumbbell-shaped guard cells flanked by subsidiary cells, whereas *A. thaliana* has kidney-shaped guard cells (Toda et al., 2016). In this study, we investigated whether *OsPATROL1* overexpression accelerates stomatal responses, thereby improving photosynthetic induction, carbon gain, and growth under fluctuating light without reducing intrinsic water-use efficiency (*iWUE)* in rice.

## Results

### OsPATROL1 is structurally similar to AtPATROL1 and is broadly expressed in rice

To characterize OsPATROL1 within the domain of unknown function 810 (DUF810) family, we compared DUF810 family proteins from *O. sativa* and *A. thaliana* (Fig. S1a). The predicted full-length structures of OsPATROL1 and AtPATROL1 showed high structural similarity (TM-score = 0.80; sequence identity = 62%). Also, the predicted MUN-like region of OsPATROL1 showed structural similarity to the experimentally determined MUN domain of rat Munc13-1 despite low sequence identity (TM-score = 0.59; sequence identity = 14%) (Fig. S1b).

Public transcriptome data (Hong et al., 2020) showed that *OsPATROL1* and *Os09g0346700* showed relatively high expression in vegetative tissues, including leaf blades and roots, compared with most other DUF810 family genes in rice, whereas *Os03g0138600* and *Os10g0471000* exhibited more restricted expression, predominantly in pollen (Fig. 1a). Histochemical analysis of β-glucuronidase (GUS) activity in *pOsPATROL1:GUS* seedlings detected promoter activity in both shoots and roots, with particularly strong signals observed in root apical meristem, lateral root primordia, the root stele, immature leaves, and the shoot meristem (Fig. 1b). GUS staining was also detected in leaf epidermal and guard cells. Using qRT-PCR, we confirmed *OsPATROL1* expression in leaf blades, leaf sheaths, unexpanded leaves, shoot bases, and roots, at approximately 5–10% of *ACTIN1* transcript abundance (Fig. 1c).

**Figure 1.**
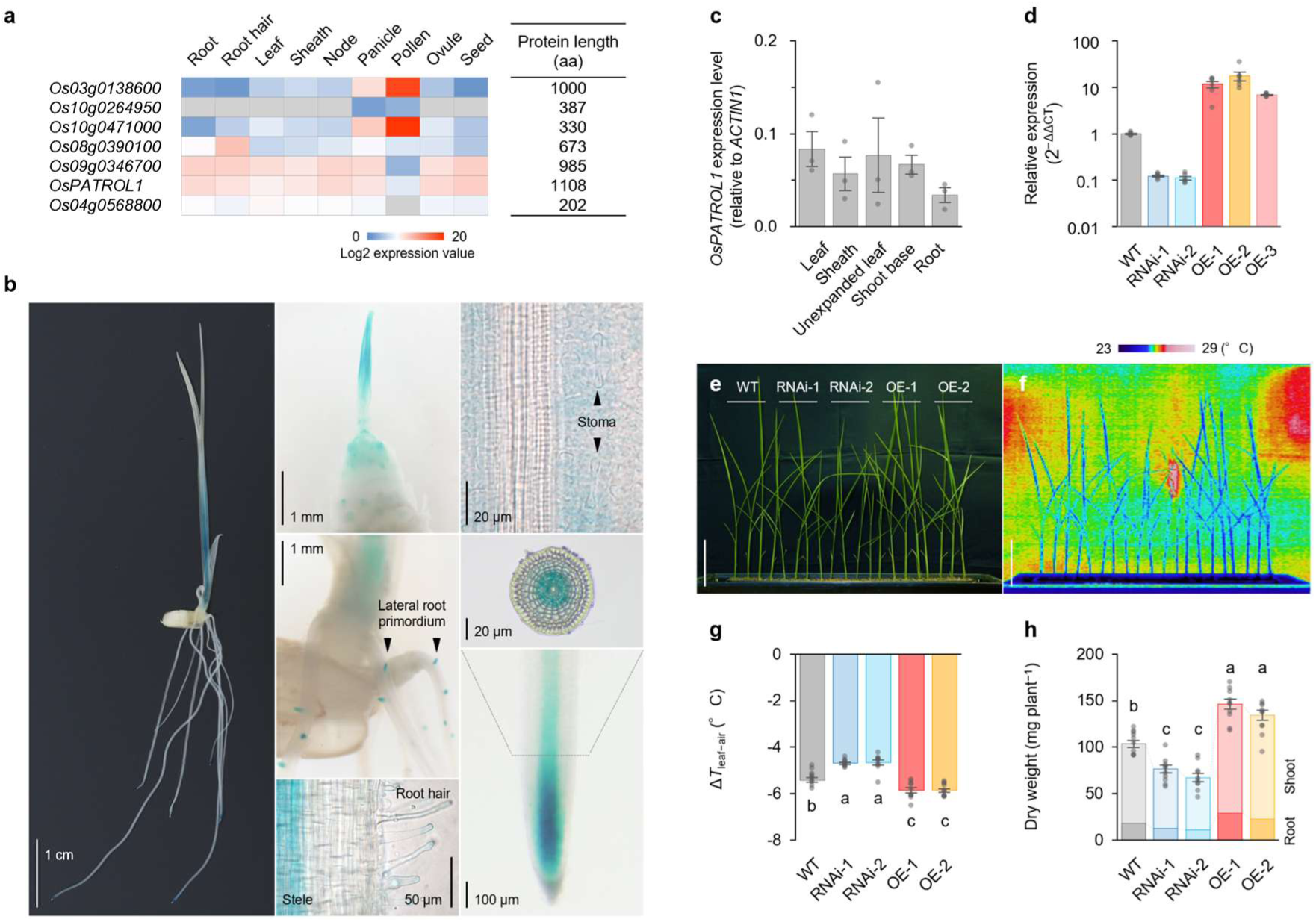
Tissue-specific expression of *OsPATROL1* and phenotypes of *OsPATROL1* knockdown and overexpression lines. (a) The tissue-specific expression profiles of *OsPATROL1* and other OsDUF810 family genes, and the lengths of their encoded proteins. Expression data were retrieved from the CAFRI-Rice database (Hong et al., 2020), and protein lengths were obtained from the RAP-DB (Kawahara et al., 2026). (b) Histochemical GUS staining of 1-week-old *pOsPATROL1:GUS* seedlings. (c) Tissue- specific expression of *OsPATROL1* in 4-week-old wild-type (WT) plants at the five-leaf stage (*n* = 3). (d) *OsPATROL1* transcript levels in the fifth leaf blade of WT, *OsPATROL1*-RNAi (RNAi), and *OsPATROL1*-overexpression (OE) lines at the five-leaf stage (*n* = 8). (e, f) Representative growth phenotypes and thermal images of WT, RNAi, and OE plants grown under glasshouse conditions at the five-leaf stage (scale bar = 10 cm). (g, h) Leaf-to-air temperature difference (Δ*T*_leaf−air_) and shoot and root dry weights of WT, RNAi, and OE plants (*n* = 10). Error bars indicate SEM. Different letters indicate significant differences among genotypes (Tukey’s test, *P* < 0.05).

To assess potential functional conservation between *AtPATROL1* and *OsPATROL1*, we compared Gene Ontology enrichment of their top 50 coexpressed genes (Obayashi et al., 2022). In both species, coexpressed genes were enriched for biological processes related to vesicle-mediated and Golgi vesicle transport and intracellular protein localization (Fig. S2, Dataset S1). These shared patterns suggest that AtPATROL1 and OsPATROL1 function in similar membrane-trafficking contexts.

To examine the physiological effects of altered *OsPATROL1* expression, we generated RNA interference (RNAi) and overexpression (OE) lines. *OsPATROL1* transcript abundance was reduced to approximately 11–12% of the wild-type (WT) levels in the two RNAi lines and increased 6.9- to 17.5-fold in the three OE lines (Fig. 1d). The transgenic lines showed corresponding differences in leaf temperature and growth at the five-leaf stage (Fig. 1e, f). The leaf-to-air temperature difference (Δ*T*_leaf−air_) was significantly higher in the RNAi lines and lower in the OE lines than in WT, consistent with reduced and enhanced transpirational cooling, respectively (Fig. 1g). Shoot and root dry weights tended to be lower in the RNAi lines and higher in the OE lines. Total dry weight was significantly lower in the RNAi lines and higher in the OE lines than in WT (Fig. 1h). Thus, *OsPATROL1* expression is associated with both transpiration and early vegetative growth.

### OsPATROL1 overexpression has limited effects on steady-state photosynthetic traits and does not alter stomatal morphology

Next, we examined whether *OsPATROL1* overexpression altered stomatal morphology or steady-state photosynthetic traits. The stomatal density and length on both the adaxial and abaxial leaf surfaces did not significantly differ among WT, OE-1, and OE-2 (Fig. 2a). Leaf nitrogen content and relative chlorophyll content per leaf area were also comparable among genotypes (Fig. 2b, c). Similarly, dark respiration rate (*R*_dark_) and nocturnal *g*_sw_ were comparable among genotypes (Fig. 2d). The response of *A* to intercellular CO₂ concentration (*C*_i_) was generally similar among genotypes, although OE-1 showed significantly higher *A* at near-ambient CO₂ concentrations and OE-2 showed a similar trend (Fig. 2e). The estimated maximum rate of carboxylation by Rubisco (*V*_cmax_) did not significantly differ among genotypes. Thus, *OsPATROL1* overexpression had modest effects on steady-state gas exchange and caused no detectable changes in stomatal morphology or major biochemical traits.

**Figure 2.**
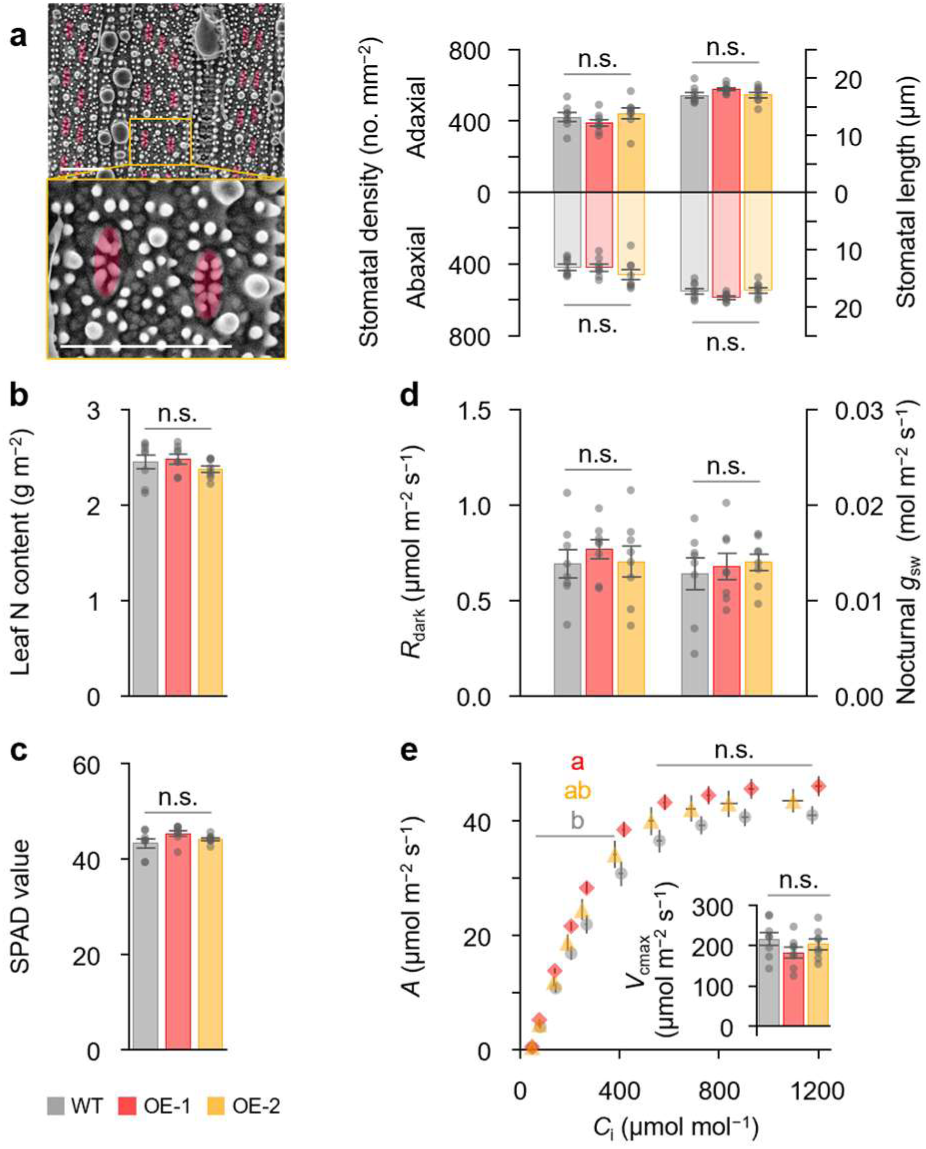
Steady-state photosynthesis of WT and *OsPATROL1*-OE plants. (a) Representative scanning electron micrographs of stomata on the abaxial leaf surface and stomatal density and length on the adaxial and abaxial leaf surfaces (scale bars = 50 μm). (b) Leaf nitrogen content. (c) Relative chlorophyll content (SPAD value). (d) Dark respiration rate (*R*_dark_) and nocturnal stomatal conductance (*g*_sw_). (e) CO₂ response of the net CO₂ assimilation rate (*A*) and estimated maximum rate of carboxylation by Rubisco (*V*_cmax_). *n* = 8. Error bars indicate SEM. Different letters indicate significant differences among genotypes (Tukey’s test, *P* < 0.05).

To test this tendency further, we measured steady-state *g*_sw_ in three independent OE lines at different irradiances in a separate experiment. At a PPFD of 150 μmol m⁻² s⁻¹, *g*_sw_ was significantly higher in OE-1 and OE-3 than in WT, whereas that in OE-2 was comparable to WT (Fig. S4). At 450 μmol m⁻² s⁻¹, no significant differences were detected, although OE-1 and OE-3 showed slightly higher *g*_sw_. At 1500 μmol m⁻² s⁻¹, all three OE lines tended to have higher *g*_sw_ than WT (*P* < 0.10) (Fig. S4). These results support a modest, condition-dependent effect of *OsPATROL1* overexpression on steady-state *g*_sw_, consistent with the slightly higher *g*_sw_ observed in *A. thaliana* (Hashimoto-Sugimoto et al., 2013).

### OsPATROL1 overexpression accelerates stomatal opening and photosynthetic induction

We next examined the gas-exchange responses to changes in irradiance. After PPFD was increased from 50 to 1500 μmol m⁻² s⁻¹, the time constant (*τ*) of *g*_sw_ during induction was significantly reduced by 43% in OE-1 and 41% in OE-2 relative to WT (Fig. 3a, b). The *τ* of *A* was similarly reduced by 44% and 46%, respectively (Fig. 3c). By contrast, *τ* values for both *g*_sw_ and *A* during relaxation after return to low light did not significantly differ among genotypes (Fig. 3b, c). Consistent with these differences in *τ*, normalized *g*_sw_ and *A* increased more rapidly in both OE lines during photosynthetic induction (Fig. 3d).

**Figure 3.**
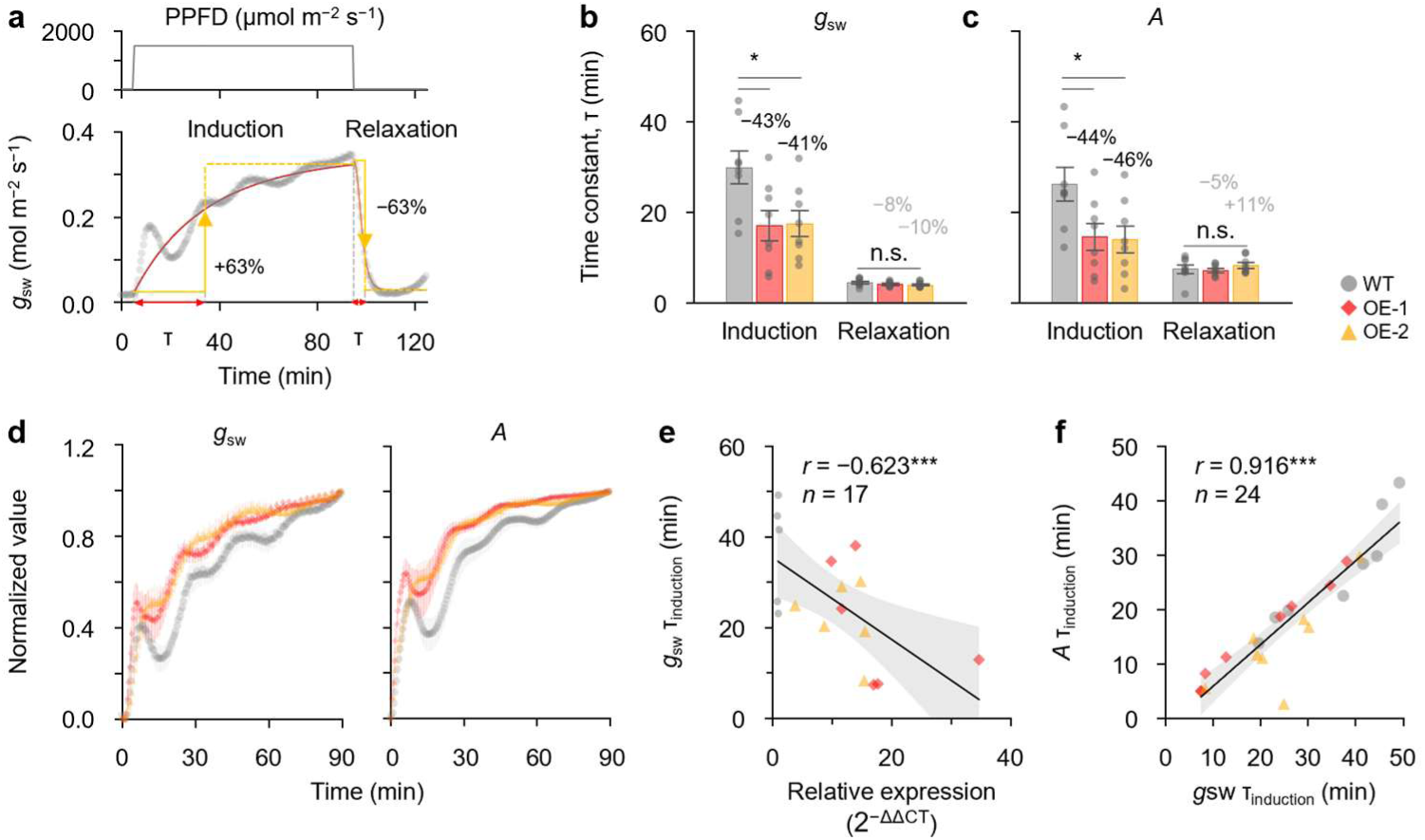
Dynamic photosynthesis of WT and *OsPATROL1*-OE plants. (a) Light regime used for photosynthetic induction and relaxation measurements and schematic definition of the time constant (τ). After overnight dark adaptation, leaves were exposed to 50 μmol m⁻² s⁻¹ PPFD for 30 min, 1500 μmol m⁻² s⁻¹ for 90 min, and then 50 μmol m⁻² s⁻¹ for 30 min. (b, c) Time constants (τ) of *g*_sw_ and *A* during induction and relaxation (*n* = 8). Error bars indicate SEM. Asterisks indicate significant differences from WT (Dunnett’s test, *P* < 0.05). Percentages indicate changes relative to WT; the black and gray percentages indicate significant and nonsignificant changes, respectively. (d) Normalized *g*_sw_ and *A* during photosynthetic induction (*n* = 8). Error bars indicate SEM. (e) Correlation between *OsPATROL1* transcript levels and τ of *g*_sw_ during induction. (f) Correlation between τ of *g*_sw_ and τ of *A* during induction. Shaded areas indicate 95% confidence intervals. *** *P* < 0.001.

Leaf *OsPATROL1* transcript abundance was negatively correlated with *τ* of *g*_sw_ during induction (*r* = −0.623, *P* < 0.001), indicating that higher *OsPATROL1* expression was associated with faster stomatal opening (Fig. 3e). Moreover, *τ* of *g*_sw_ was strongly correlated with *τ* of *A* (*r* = 0.916, *P* < 0.001), supporting a close association between faster stomatal opening and photosynthetic induction (Fig. 3f).

During induction, the time-averaged *g*_sw_ was 48% and 16% higher in OE-1 and OE-2, respectively, than in WT (Fig. S3a). *A* was also higher in both OE lines during induction, including under low light (Fig. S3b). The *iWUE* was not reduced in either OE line during induction and relaxation (Fig. S3c). The electron transport rate through photosystem II (ETRII) tended to be higher during induction, with mean values 23% and 12% higher in OE-1 and OE-2, respectively; in contrast, no clear differences were observed during relaxation (Fig. S3d). Thus, *OsPATROL1* overexpression accelerates stomatal opening and photosynthetic induction without compromising *iWUE*.

### OsPATROL1 overexpression alleviates growth reductions under fluctuating light

To examine whether accelerated photosynthetic induction affected plant growth under fluctuating light, we grew the WT and three independent OE lines under steady- or fluctuating-light with the same daily integrated PPFD (Fig. 4a). Through a two-way ANOVA, we showed significant effects of genotype, light treatment, and their interaction on tiller number and shoot dry weight (Fig. 4b, d).

**Figure 4.**
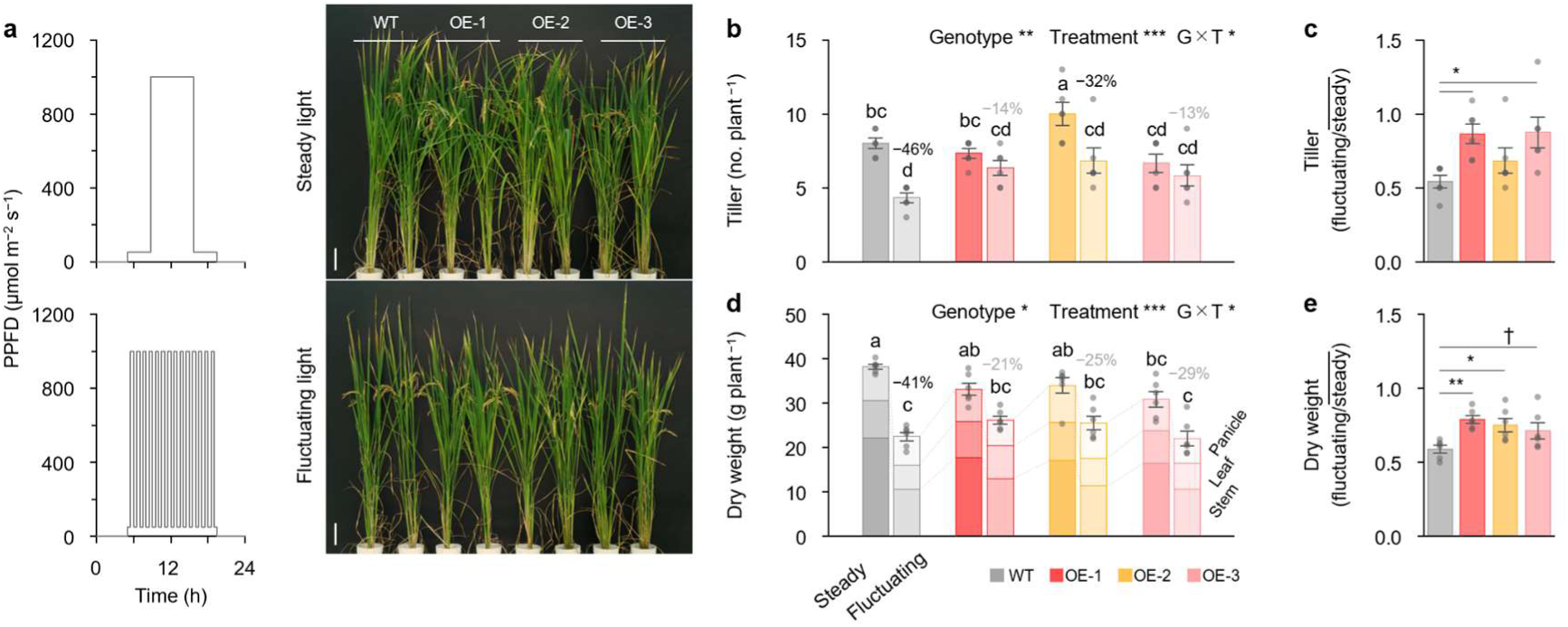
Growth of WT and *OsPATROL1*-OE plants under artificial fluctuating light. (a) Steady- and fluctuating-light treatments with equivalent daily integrated PPFD and representative growth phenotypes under each light regime (scale bar = 10 cm). (b, d) The tiller number and shoot dry weight under steady and fluctuating light (*n* = 6). Error bars indicate SEM. Effects of genotype, light treatment, and their interaction were tested by two-way ANOVA. Different letters indicate significant differences (Tukey’s test, *P* < 0.05). Percentages indicate changes under fluctuating light relative to steady light; the black and gray percentages indicate significant and nonsignificant changes, respectively. (c, e) Tiller number and shoot dry weight under fluctuating light relative to steady light (Dunnett’s test vs WT: ** *P* < 0.01, * *P* < 0.05, † *P* < 0.10).

Tiller number decreased by 46% in WT under fluctuating light relative to steady light, whereas the reductions were smaller in OE-1 (14%), OE-2 (32%), and OE-3 (13%) (Fig. 4b, c). Shoot dry weight was similarly reduced by 41% in WT but only 21%, 25%, and 29% in OE-1, OE-2, and OE-3, respectively (Fig. 4d). Consistent with the significant genotype × light interaction, the relative reduction in shoot dry weight under fluctuating light was smaller in the OE lines than in WT (Fig. 4e). These results indicate that *OsPATROL1* overexpression alleviates the growth reduction caused by fluctuating light.

### Faster stomatal responses enhance carbon gain under simulated natural fluctuating light without reducing iWUE

To determine whether the faster stomatal responses of the OE lines enhanced photosynthesis under a field-relevant pattern of light fluctuations, we measured gas exchange for 12 h using a PPFD pattern recorded in the middle canopy of field-grown rice (Fig. 5a). Throughout the fluctuating-light period, *g*_sw_ and *A* were generally higher in OE-1 and OE-2 than in WT (Fig. 5b, c). The 12-h mean *A* was 12% higher in OE-1 and 8% higher in OE-2 than in WT (Fig. 5c). Despite the higher *g*_sw_, *iWUE* was not reduced in either OE line (Fig. 5d).

**Figure 5.**
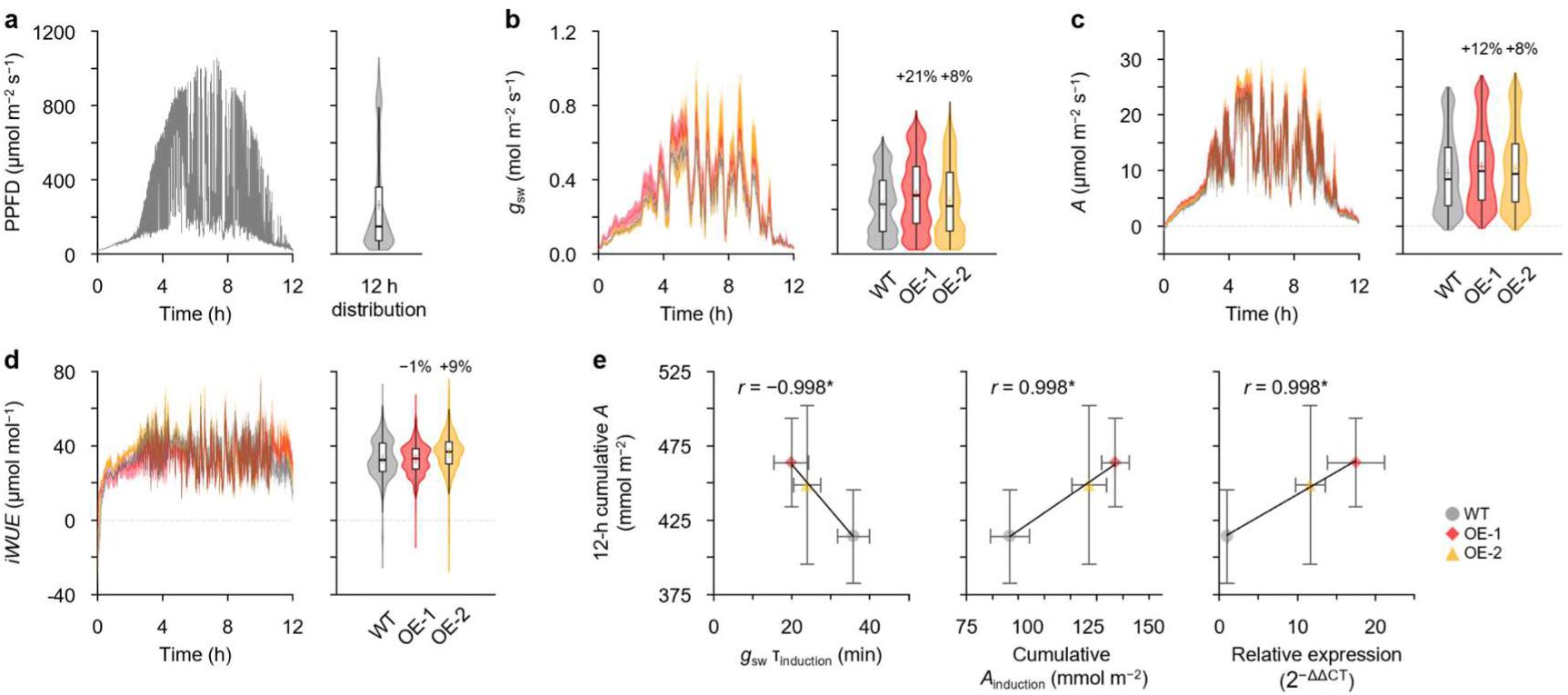
Photosynthetic performance under simulated natural fluctuating light. (a) Diurnal PPFD pattern recorded in the middle canopy of field-grown rice and its 12 h distribution. (b–d) Diurnal changes and distributions of *g*_sw_, *A*, and intrinsic water-use efficiency (*iWUE*) in WT and *OsPATROL1*-OE plants during 12 h of simulated natural fluctuating light (*n* = 4–5). Error bars indicate SEM. Percentages indicate changes relative to WT. (e) Correlations of genotype means (*n* = 3) for 12-h cumulative *A* with τ of *g*_sw_ during induction, cumulative *A* during a single induction, and leaf *OsPATROL1* transcript levels. * indicates *P* < 0.05.

Across the three genotypes, the 12-h cumulative *A* was negatively associated with *τ* of *g*_sw_ during induction (Fig. 5e). The 12-h cumulative *A* was also positively associated with cumulative *A* during a single induction event and with leaf *OsPATROL1* transcript abundance (Fig. 5e). These relationships are consistent with the accelerated stomatal and photosynthetic induction observed during a single light transition contributing to greater carbon gain under simulated natural fluctuating light.

### OsPATROL1 overexpression enhances vegetative growth under glasshouse conditions

We next examined whole-plant growth under glasshouse conditions. Compared with WT, both OE lines showed greater vegetative growth (Fig. 6a), with increased tiller number and leaf area (Fig. 6b, c). Relative to that in WT, total dry weight was significantly increased by 34% in OE-1 and 44% in OE-2 (Fig. 6d). The bleeding sap rate was also higher in both OE lines (Fig. 6e), accompanied by increased leaf nitrogen content (Fig. 6f).

**Figure 6.**
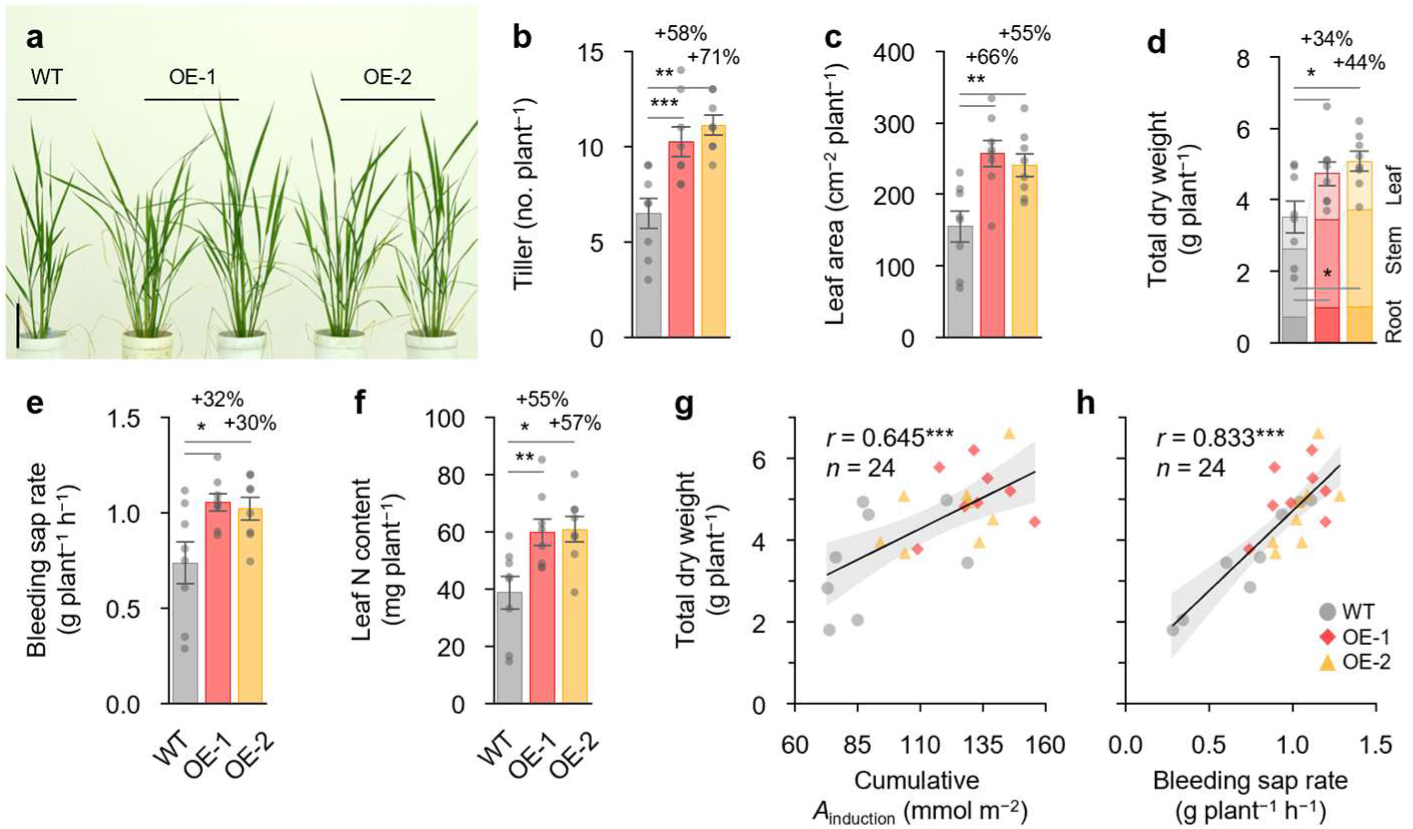
Growth and resource acquisition of WT and *OsPATROL1*-OE plants under glasshouse conditions. (a) Representative growth phenotypes of WT and *OsPATROL1*-OE plants (scale bar = 10 cm). (b–f) Tiller number, leaf area, total dry weight, bleeding sap rate, and leaf nitrogen content (*n* = 8). Error bars indicate SEM. Asterisks indicate significant differences from WT (Dunnett’s test; *** *P* < 0.001, ** *P* < 0.01, * *P* < 0.05). Percentages indicate changes relative to WT. (g, h) Relationships of total dry weight with cumulative *A* during photosynthetic induction and bleeding sap rate, respectively (*n* = 24). Shaded areas indicate 95% confidence intervals. *** *P* < 0.001.

Across individual plants, the total dry weight was positively correlated with cumulative *A* during photosynthetic induction (*r* = 0.645, *P* < 0.001; Fig. 6g) and with bleeding sap rate (*r* = 0.833, *P* < 0.001; Fig. 6h). Additional analyses showed positive relationships among tiller number, root dry weight, bleeding sap rate, leaf nitrogen content, and leaf carbon content (Fig. S5). Taken together, the enhanced vegetative growth was accompanied by greater photosynthetic carbon gain, root biomass, bleeding sap rate, and leaf nitrogen status.

We also assessed reproductive performance under pot-grown glasshouse conditions over two years (Fig. S6). Although the OE lines generally produced 19–55% more panicles, the grain yield was not consistently increased across lines and years because increases in panicle number were partly offset by lower single-grain weight in some lines (Fig. S6g–j). Thus, the vegetative growth advantage observed here did not translate consistently into greater grain yield under the pot conditions tested.

## Discussion

### OsPATROL1 accelerates stomatal opening and dynamic photosynthesis without reducing iWUE

Although steady-state *g*_sw_ and *A* tended to be higher in the OE lines, these effects varied among the lines and measurement conditions, whereas the induction kinetics were consistently enhanced: *τ* of *g*_sw_ was reduced by 41–43% and *τ* of *A* by 44–46% in both OE lines (Fig. 3). The stomatal density and length, leaf nitrogen and relative chlorophyll contents, *R*_dark_, nocturnal *g*_sw_, and *V*_cmax_ were unchanged (Fig. 2), indicating that faster photosynthetic induction occurred without detectable changes in stomatal morphology or photosynthetic biochemical capacity. Moreover, *τ* of *g*_sw_ was strongly correlated with *τ* of *A* (*r* = 0.916), indicating that faster stomatal opening contributed to enhanced photosynthetic induction. ETRII also tended to be higher during induction but not during relaxation (Fig. S3), possibly because faster stomatal opening alleviated CO₂ limitation following the increase in irradiance, whereas electron transport remained light-limited after the return to low irradiance (Adachi et al., 2019; Yamori et al., 2020).

Previous studies in rice have shown that increasing *g*_sw_ through loss of *SLAC1* or *OSA1* overexpression can enhance *A* but may increase water loss or reduce water-use efficiency (Kusumi et al., 2012; Yamori et al., 2020; Zhang et al., 2021). The dynamic phenotype of *OsPATROL1*-OE plants was distinct from a constitutively open phenotype. Although steady-state *g*_sw_ tended to be higher under some conditions (Figs. S3, S4), the more consistent effect was faster opening after increasing irradiance, whereas the relaxation kinetics were unchanged (Fig. 3). Under the 12-h simulated natural fluctuating light, this temporal advantage increased the cumulative *A* by 8–12% while maintaining *iWUE* (Fig. 5). Across the three genotypes, a smaller *τ* of *g*_sw_ was associated with a greater 12-h cumulative *A* (Fig. 5e), consistent with the kinetic advantage measured during a single light transition contributing to daytime carbon gain. A similar phenotype occurs in *PATROL1*-overexpressing *A. thaliana*, in which faster stomatal opening enhances photosynthetic induction and biomass without reducing the water-use efficiency (Kimura et al., 2020). Thus, unlike approaches that constitutively increase stomatal conductance, *OsPATROL1* accelerates stomatal opening when the photosynthetic demand increases.

### OsPATROL1 may share a conserved membrane-trafficking function

The predicted structures of OsPATROL1 and AtPATROL1 were highly similar (Fig. S1). In addition, *OsPATROL1* promoter activity was detected in rice guard cells, and genes coexpressed with *PATROL1* in both species were enriched for vesicle- and Golgi-mediated transport (Figs. 1, S2).

In *A. thaliana*, PATROL1 regulates the stimulus-dependent translocation of the plasma membrane H⁺-ATPase AHA1 in guard cells (Hashimoto-Sugimoto et al., 2013) and was identified in plant SNARE interactomes (Fujiwara et al., 2014). Considering that the MUN domain of Munc13-1 facilitates vesicle priming by promoting SNARE-complex assembly (Ma et al., 2011), OsPATROL1-dependent recruitment of H⁺-ATPases to the plasma membrane could explain why stomatal opening was accelerated while the relaxation kinetics remained unchanged (Fig. 3). The CO₂-dependent redistribution of the K⁺ channel KAT1 provides a related example of transporter trafficking regulating stomatal kinetics without changes in the expression of KAT1 itself (Yu et al., 2026). Future studies should therefore determine whether OsPATROL1 directly regulates H⁺- ATPase trafficking in rice.

### Enhanced carbon gain is associated with greater vegetative growth

The dynamic-photosynthesis phenotype of *OsPATROL1*-OE plants was accompanied by improved vegetative growth. Under artificial fluctuating light with the same daily integrated PPFD as the steady-light treatment, shoot dry weight decreased by 41% in WT but by only 21–29% in the OE lines. Furthermore, only WT showed a significant reduction in shoot dry weight between light regimes (Fig. 4). Nevertheless, shoot dry weight under artificial fluctuating light did not differ significantly among genotypes, possibly because higher *g*_sw_ increased water loss from the OE lines under low humidity in the controlled-environment room (approximately 50%), limiting their growth advantage. Under glasshouse conditions, the total dry weight was 34–44% greater in the OE lines and was positively associated with cumulative *A* during photosynthetic induction (Fig. 6).

In addition, *OsPATROL1*-OE plants showed greater tiller number, root biomass, bleeding sap rate, and leaf nitrogen content (Fig. 6; Fig. S5). The increase in tiller number may result from enhanced carbon gain, as reduced photosynthesis suppresses tillering, whereas increased sugar availability promotes tiller-bud outgrowth in rice (Yamamoto et al., 1995; Wen et al., 2024). PATROL1 functions in the roots of *A. thaliana*, where it contributes to root meristem activity, root elongation and branching, and whole-plant growth (Notaguchi et al., 2024; Katsuhama et al., 2025). These root-associated traits may contribute to the growth phenotype, although the present data do not distinguish whether they are a cause or a consequence of enhanced whole-plant growth.

Enhanced vegetative growth did not consistently translate into greater grain yield under the pot-grown glasshouse conditions used in this study (Fig. S6). Increased tiller and panicle numbers were partly offset by reductions in single-grain weight in some OE lines. The root size limitation and associated constraints on resource availability under pot conditions may have reduced *A* and growth later in development, thereby constraining expression of the photosynthetic advantage of the OE lines (Poorter et al., 2012). Greater tillering may also have increased mutual shading within the canopy, reducing light availability and *A* later during development (Navasero and Tanaka, 1966). Field experiments will therefore be necessary to determine whether the dynamic photosynthetic and vegetative-growth advantages of *OsPATROL1* overexpression are maintained in agronomic environments.

Taken together, *OsPATROL1* overexpression accelerates stomatal opening and improves photosynthetic induction and daytime carbon gain under fluctuating light conditions without reducing *iWUE*. The associated mitigation of fluctuating-light growth penalties identifies stomatal kinetics as a promising target for improving crop carbon gain and vegetative performance under dynamic light environments.

## Materials and methods

### Plant materials and generation of transgenic rice lines

*Oryza sativa* L. cv. Nipponbare was used as the wild type (WT) and as the genetic background for all transgenic lines. To generate *OsPATROL1*-overexpressing lines (*OsPATROL1*-OE), we isolated total RNA from Nipponbare leaves using the RNeasy Plant Mini Kit (Qiagen, Germany), and first-strand cDNA was synthesized using oligo(dT)₁₈ primers and the PrimeScript II First-Strand cDNA Synthesis Kit (Takara, Japan). We amplified the full-length coding sequence of *OsPATROL1* by RT-PCR using gene-specific primers (Table S1) and cloned it into the binary vector pBI-Hm under the control of the rice *Actin* promoter.

To generate *OsPATROL1* knockdown lines (*OsPATROL1*-RNAi), we amplified a 363-bp fragment of the 3′-UTR of *OsPATROL1* from cDNA using gene-specific primers (Table S1) and introduced it into the RNA interference vector pANDA (Miki and Shimamoto, 2004). For *OsPATROL1* promoter:GUS reporter lines (*pOsPATROL1:GUS*), a 2.2-kb genomic region comprising the *OsPATROL1* promoter and the 5′-UTR was obtained from Nipponbare genomic DNA and cloned upstream of a promoterless *GUS* gene in the binary vector pBI101-Hm using the NEBuilder HiFi DNA Assembly Kit (New England Biolabs, USA).

The resulting binary constructs were introduced into Nipponbare calli by *Agrobacterium tumefaciens*-mediated transformation as described previously (Hiei et al., 1994). Transformed calli were selected on hygromycin-containing medium and regenerated into whole plants. Independent *OsPATROL1*-OE and *OsPATROL1*-RNAi lines were screened based on *OsPATROL1* transcript abundance in leaves determined by qRT-PCR and used for subsequent analyses.

### Growth conditions

Seeds were germinated in cell trays, transferred to 1 L (2024) or 3 L (2025) pots, and grown in a glasshouse at the University of Tokyo, Japan (35°43′00″N, 139°45′45″E). Pots were filled with a commercial sandy loam soil for rice cultivation (Honens Soil No. 1; HONENAGRI Co., Ltd., Japan). Before transplanting, a slow-release fertilizer was incorporated at 0.1 g L⁻¹ N, P, and K, in addition to the inherent soil nutrient content (0.6 g L⁻¹ N, P, and K). Pots were arranged at 20-cm intervals in water-filled trays, and intermittent irrigation was applied throughout the growth period. To minimize any positional effects within the glasshouse, we randomly repositioned all pots each week. During the growth period, the mean ± SD of daily solar radiation, daily mean air temperature, relative humidity, and air vapour pressure deficit were 15.5 ± 6.9 MJ m^−2^ day^−1^, 27.8 ± 3.9°C, 75.5 ± 8.6%, and 1.16 ± 0.58 kPa in 2024, and 18.0 ± 7.1 MJ m^−2^ day^−1^, 28.1 ± 4.0°C, 74.6 ± 8.8%, and 1.24 ± 0.56 kPa in 2025, respectively.

### Artificial fluctuating light experiment

Plants were grown in a controlled-environment room maintained at 28°C. Thirty-day-old seedlings were transplanted into 1-L pots and acclimated for an additional 15 days under steady light. Thereafter, the plants were subjected to either steady-light or fluctuating-light conditions for 120 days. In the steady-light treatment, PPFD was set to 50 μmol m⁻² s⁻¹ during the first and last 3.5 h of the 14- h photoperiod and 1000 μmol m⁻² s⁻¹ during the middle 7 h. For the fluctuating-light treatment, the PPFD was alternated between 50 and 1000 μmol m⁻² s⁻¹ at 30-min intervals, and the daily integrated PPFD was matched to that of the steady-light treatment.

### Histochemical GUS staining

One-week-old *pOsPATROL1:GUS* plants grown vertically on solid medium (1/2 MS, 1% sucrose, 0.05% 2-(*N*-morpholino)ethanesulfonic acid (MES), 0.1% Plant Preservative Mixture (PPM; Plant Cell Technology, USA), pH 5.7, 0.8% gellan gum) were prefixed in 90% acetone at −20°C for 1 h. The fixed samples were rinsed several times in 100 mM sodium phosphate buffer (pH 7.0) and incubated in GUS staining buffer (100 mM sodium phosphate buffer [pH 7.0], 0.1% Triton X-100, 5 mM potassium ferrocyanide, 5 mM potassium ferricyanide, 0.5 mg/mL 5-bromo-4-chloro-3-indolyl-β-D-glucuronic acid, 10% methanol) for 12–24 h at 37°C. The stained samples were rinsed several times with 70% ethanol, transferred to ethanol/acetic acid (6:1, v/v) for 72 h, and finally cleared in chloral hydrate/water/glycerol (8:1:2, w/v/v) before observation.

### qRT-PCR

For tissue-specific expression analysis, leaf blades and sheaths were collected from the fully expanded fifth leaf, unexpanded leaves from the sixth leaf, shoot bases from the basal 5 mm of the shoot, and roots from 4-week-old WT plants at the five-leaf stage. Total RNA was extracted using the RNeasy Plant Mini Kit (QIAGEN, Germany), and cDNA was synthesized using the PrimeScript FAST RT Reagent Kit with gDNA Eraser (Takara Bio, Japan). qRT-PCR was performed using TB Green Premix Ex Taq II (Tli RNase H Plus) (Takara Bio, Japan). Tissue-specific expression relative to *ACTIN1* was calculated using the 2^−ΔCT^ method, whereas expression in transgenic lines relative to WT was calculated using the 2^−ΔΔCT^ method. The amplification efficiencies of the *OsPATROL1* and *ACTIN1* primer pairs were 100.36% and 98.61%, respectively. The primer sequences are listed in Table S1.

### Thermal imaging

Thermal images were acquired using a far-infrared camera (Thermo Tracer TH9100MR; NEC Avio Infrared Technologies, Japan) in the glasshouse on a clear morning. Leaf-to-air temperature differences (Δ*T*_leaf−air_) were calculated by subtracting air temperature from leaf temperature.

### Leaf gas exchange and chlorophyll fluorescence

Measurements were conducted on the central portion of the second-youngest fully expanded leaves between panicle initiation and booting (74–81 days after planting). The photosynthetic responses to step changes in light intensity were measured using an LI-6800 portable photosynthesis system (LI-COR, USA). After overnight dark adaptation, leaves were enclosed in the LI-6800 chamber in darkness for 10 min, and dark respiration rate (*R*_dark_) and nocturnal stomatal conductance (*g*_sw_) were measured. Leaves were then acclimated for 30 min at a PPFD of 50 μmol m⁻² s⁻¹, air temperature of 28°C, relative humidity of 65%, and reference CO₂ concentration of 400 μmol mol⁻¹. The PPFD was then increased to 1500 μmol m⁻² s⁻¹ for 90 min and subsequently decreased to 50 μmol m⁻² s⁻¹ for 30 min. The net CO₂ assimilation rate (*A*) and *g*_sw_ were recorded every 30 s. Time constants (τ) for the induction and relaxation of *g*_sw_ and *A* were calculated as described previously (Vialet-Chabrand et al., 2017), where τ represents the time required to reach 63% of the total change. Normalized *g*_sw_ and *A* were calculated as (*x* − initial)/(final − initial). Intrinsic water-use efficiency (*iWUE*) was calculated as *A*/*g*_sw_. The cumulative *A* during the 90-min induction period was calculated by summing *A* multiplied by the 30-s measurement interval. Electron transport rate through PSII (ETRII) was calculated as Y(II) × PPFD × 0.5 × 0.84, where 0.5 represents the fraction of absorbed light partitioned to PSII and 0.84 represents leaf absorptance.

After the step-change measurements, leaves were acclimated at a PPFD of 1500 μmol m⁻² s⁻¹ for 30 min, and *A* was measured at reference CO₂ concentrations of 400, 50, 100, 200, 300, 400, 600, 800, 1000, 1200, and 1500 μmol mol⁻¹. The maximum rate of carboxylation by Rubisco (*V*_cmax_) was estimated by fitting *A/C*_i_ curves as described previously (Sharkey, 2016).

The steady-state *g*_sw_ at PPFDs of 150, 450, and 1500 μmol m⁻² s⁻¹ was measured using an LI-600 porometer/fluorometer (LI-COR, USA) after *g*_sw_ had stabilized.

Diurnal changes in *g*_sw_ and *A* under simulated natural fluctuating light were measured using an LI-6400XT portable photosynthesis system (LI-COR, USA). After overnight dark adaptation, the leaves were exposed for 12 h to the fluctuating-light pattern recorded in the middle canopy of a rice paddy, as described previously (Kimura et al., 2020). The measurements were recorded every 10 s. Air temperature, relative humidity, and reference CO₂ concentration were maintained as described above. Twelve-hour cumulative *A* was calculated by summing *A* multiplied by the 10-s measurement interval.

### Leaf total nitrogen and chlorophyll contents

The leaf total nitrogen content was measured in the same leaves used for gas-exchange measurements using an NC analyzer (SUMIGRAPH NC-22F; Sumika Chemical Analysis Service, Japan). Relative chlorophyll content was measured nondestructively using a SPAD-502 chlorophyll meter (Konica Minolta, Japan).

### Stomatal traits

The adaxial and abaxial surfaces of fresh leaf samples from the central portion of the same leaves used for gas-exchange measurements were directly observed using a scanning electron microscope (JCM-6000; JEOL, Japan). More than 40 stomatal complexes from 2–3 fields of view were analyzed per biological replicate. The stomatal density and length were quantified using a machine-learning model trained on manually annotated images in Biodock (www.biodock.ai) (Xie et al., 2021).

### Bleeding sap rate

Bleeding sap rate, reflecting root xylem sap exudation driven by root pressure, was measured shortly after dawn in 80-day-old plants as described previously (Katsuhama et al., 2026). Shoots were cut 10 cm above the soil surface, and the cut ends were covered with pre-weighed cotton and wrapped with Parafilm M (Amcor, USA). After 1 h, the cotton was reweighed, and the increase in mass was used to calculate the bleeding sap rate.

### Bioinformatic analyses

Amino acid sequences of DUF810 family proteins from *A. thaliana* and *O. sativa* were aligned using MUSCLE, and a phylogenetic tree was constructed using the maximum likelihood method in MEGA version 12.1 with 1000 bootstrap replicates (Kumar et al., 2024). The AlphaFold-predicted structures of AtPATROL1 and OsPATROL1 and experimentally determined MUN domain of rat Munc13-1 (PDB 5UF7) were compared using TM-align implemented in the RCSB PDB Pairwise Structure Alignment tool (Bittrich et al., 2024).

The top 50 genes coexpressed with *AtPATROL1* and *OsPATROL1* were retrieved separately from ATTED-II (Obayashi et al., 2022) and are provided in Supplementary Dataset S1. Gene Ontology biological process enrichment analysis was performed using ShinyGO (Ge et al., 2020).

### Statistical analysis

Data were analyzed by one- or two-way ANOVA followed by Tukey’s test or Dunnett’s test, as appropriate, using R version 4.4.0 (https://www.r-project.org/). Sample sizes and statistical tests are indicated in the figure legends.

## Acknowledgements

We thank Dr Koichi Morita (Kobe University) for generating the *OsPATROL1* overexpression lines, the late Dr Ko Shimamoto (Nara Institute of Science and Technology) for generously providing the pANDA vector, and Manami Taura (Ochanomizu University) for assistance with vibratome sectioning.

## Author contributions

N.K. and W.Y. conceived the study. N.K. performed the experiments, analyzed the data, acquired funding, and wrote and revised the manuscript. R.M. generated the *OsPATROL1*-RNAi lines and contributed to rice cultivation. Y.W. generated the *pOsPATROL1:GUS* reporter lines, and S.P. contributed to rice cultivation. H.F. contributed to the generation of the *OsPATROL1*-RNAi lines and provided methodological advice. N.A. provided research facilities and scientific advice. I.T. and W.Y. supervised the study, contributed to data interpretation, acquired funding, and reviewed and edited the manuscript.

## Data availability

All data supporting the findings of this study are included in the main figures and supplementary data.

---

**Supplementary Figure S1.**
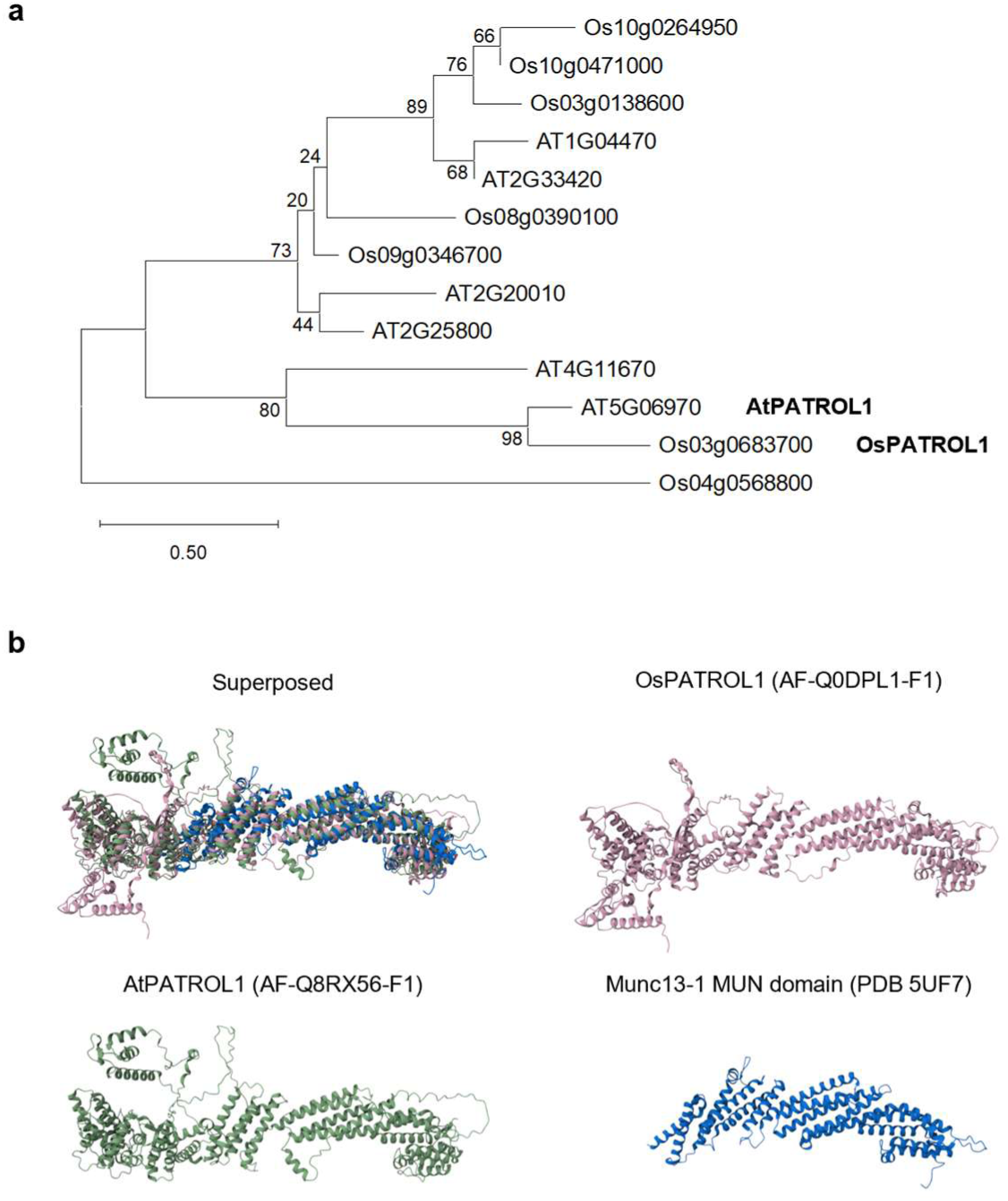
Phylogenetic relationship and structural comparison of OsPATROL1. (a) Phylogenetic tree of DUF810 family proteins from *Arabidopsis thaliana* and *Oryza sativa*. The numbers at nodes indicate bootstrap values. (b) Structural comparison of the AlphaFold-predicted full-length structures of AtPATROL1 and OsPATROL1 with the experimentally determined MUN domain of rat Munc13-1 (PDB 5UF7). Full-length OsPATROL1 and AtPATROL1 showed high structural similarity (TM-score = 0.80; sequence identity = 62%). The predicted MUN-like region of OsPATROL1 (residues 479–1103) also showed structural similarity to the Munc13-1 MUN domain despite the low sequence identity (TM-score = 0.59; sequence identity = 14%).

**Supplementary Figure S2.**
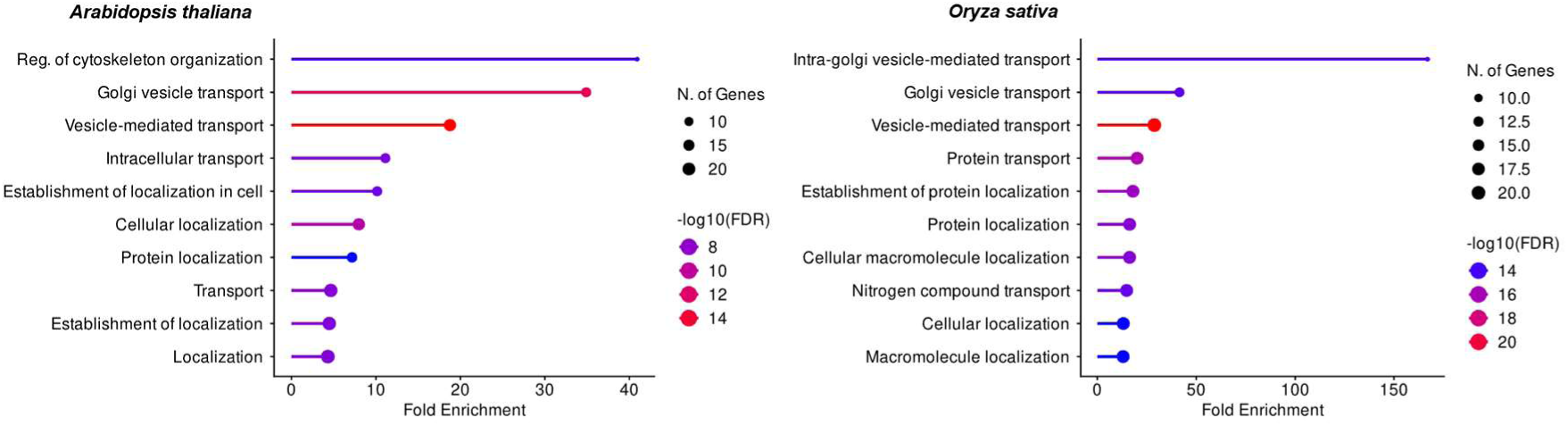
Gene Ontology enrichment analysis of genes coexpressed with *AtPATROL1* and *OsPATROL1*. Gene Ontology biological process enrichment analysis of the top 50 genes coexpressed with *AtPATROL1* in *Arabidopsis thaliana* and the top 50 genes coexpressed with *OsPATROL1* in *Oryza sativa*.

**Supplementary Figure S3.**
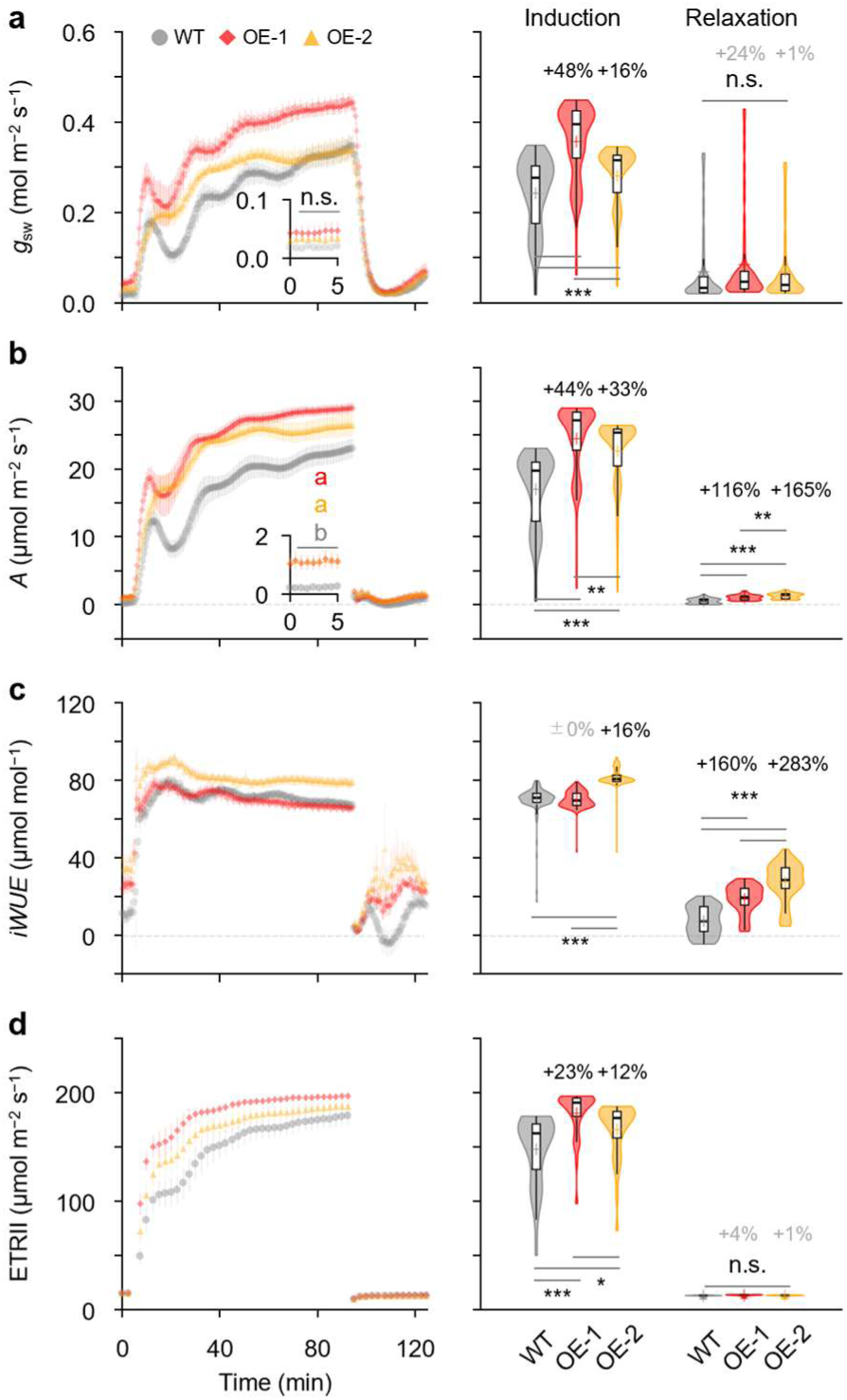
Photosynthetic responses of WT and *OsPATROL1*-OE plants during light transitions. (a–d) Temporal changes and distributions of *g*_sw_, *A*, *iWUE*, and electron transport rate through PSII (ETRII), respectively, during photosynthetic induction and relaxation (*n* = 8). Error bars indicate SEM. Percentages indicate changes relative to WT; the black and gray percentages indicate significant and nonsignificant changes, respectively. Different letters indicate significant differences among genotypes (Tukey’s test, *P* < 0.05). Asterisks indicate significant differences from WT (Tukey’s test; *** *P* < 0.001, ** *P* < 0.01, * *P* < 0.05).

**Supplementary Figure S4.**
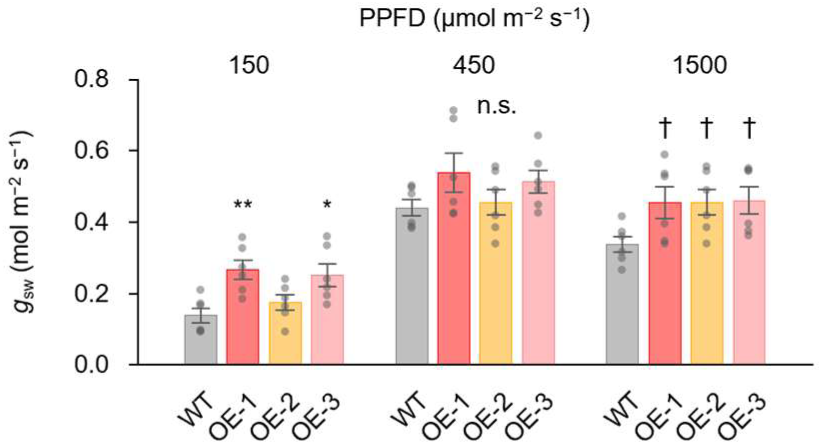
Stomatal conductance of WT and *OsPATROL1*-OE plants under different light intensities. *g*_sw_ at PPFDs of 150, 450, and 1500 μmol m⁻² s⁻¹ (*n* = 6). Error bars indicate SEM. Dunnett’s test vs WT: ** *P* < 0.01, * *P* < 0.05, † *P* < 0.10.

**Supplementary Figure S5.**
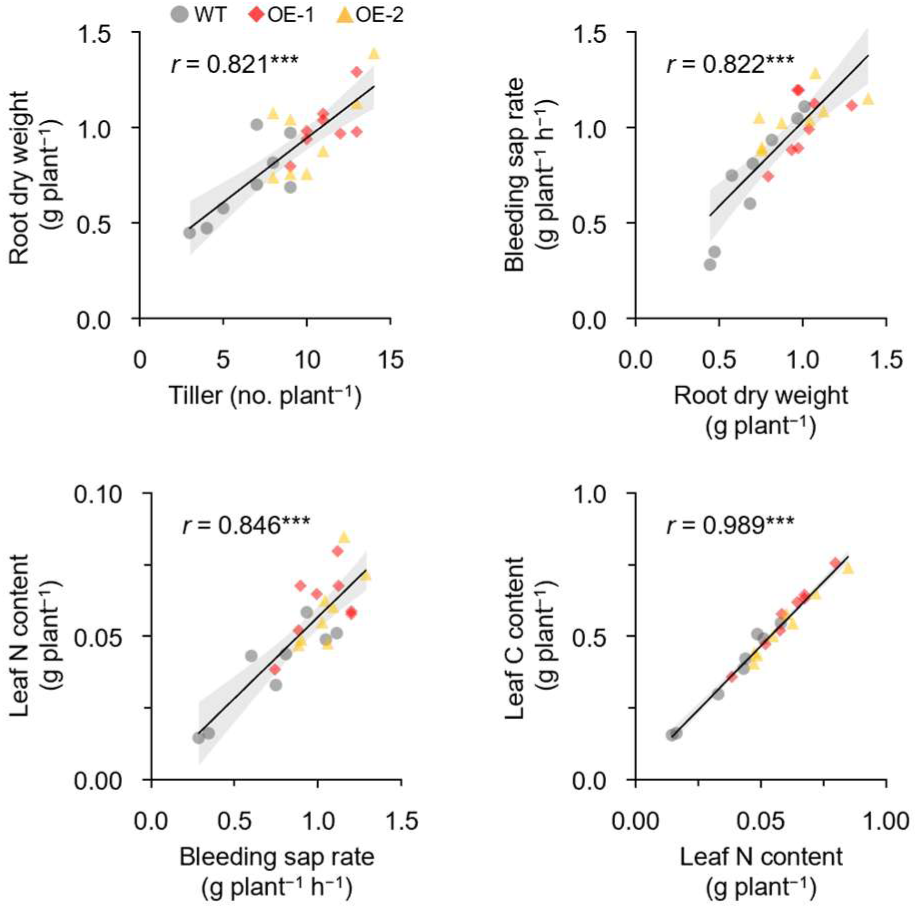
Relationships among growth and resource-acquisition traits in WT and *OsPATROL1*-OE plants. Relationships among tiller number, root dry weight, bleeding sap rate, leaf nitrogen and carbon contents (*n* = 24). The shaded areas indicate 95% confidence intervals. *** *P* < 0.001.

**Supplementary Figure S6.**
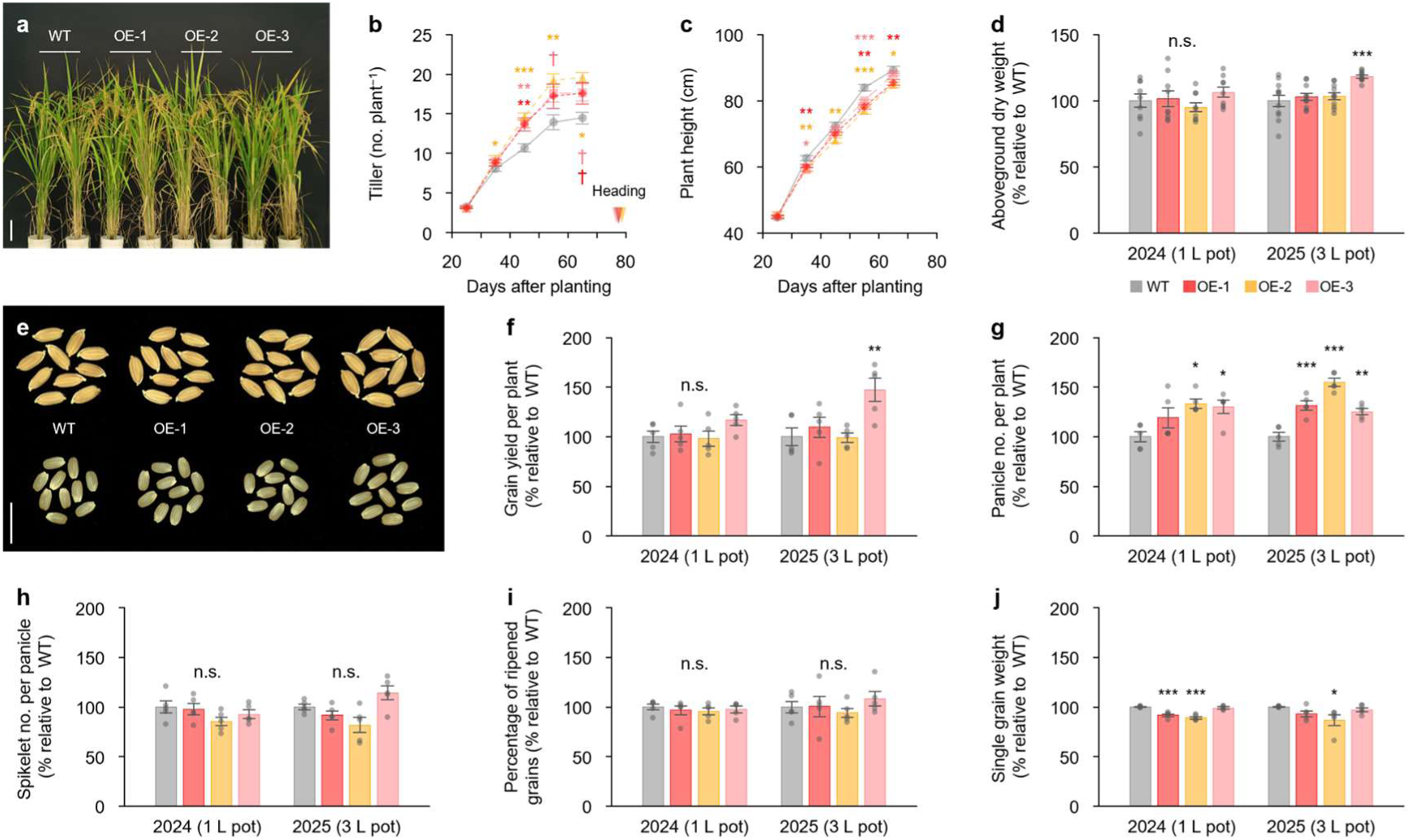
Growth and two-year yield components of WT and *OsPATROL1*-OE plants under glasshouse conditions. (a) Representative growth phenotypes of WT and *OsPATROL1*-OE plants in 2024 (scale bar = 10 cm). (b, c) Temporal changes in tiller number and plant height in 2024 and heading dates (*n* = 9). (d) The total aboveground dry weight at harvest relative to WT in plants grown in 1-L pots in 2024 (*n* = 9) and 3-L pots in 2025 (*n* = 11). (e) Representative images of harvested grains and brown rice in 2024 (scale bar = 1 cm). (f–j) Grain yield per plant and yield components relative to WT in 2024 and 2025 (*n* = 5). Error bars indicate SEM. Dunnett’s test vs WT: *** *P* < 0.001, ** *P* < 0.01, * *P* < 0.05, † *P* < 0.10.

**Supplementary Table S1.**
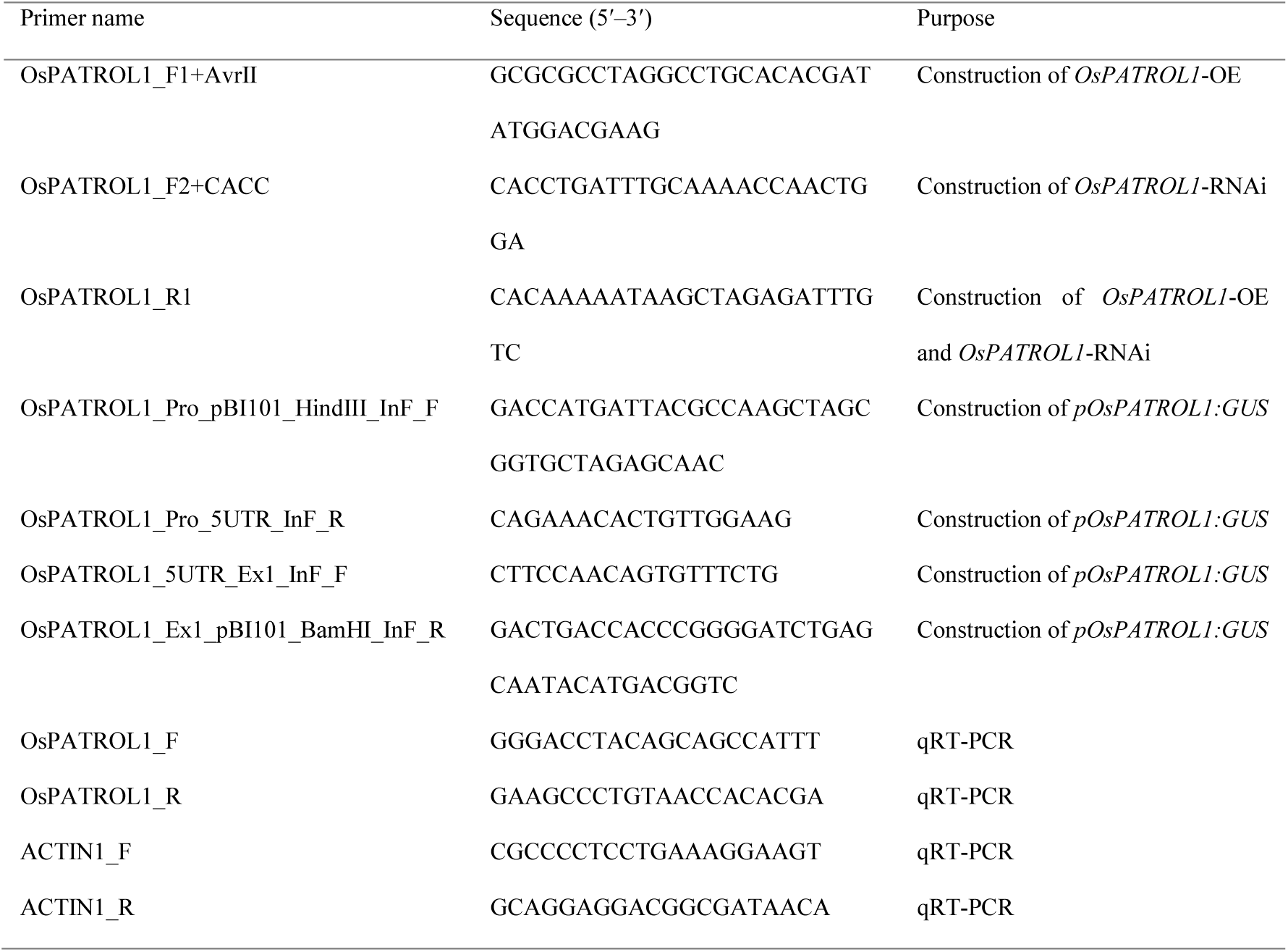
List of primers used in this study. The amplification efficiencies of the *OsPATROL1* and *ACTIN1* primer pairs for qRT-PCR were 100.36% and 98.61%, respectively.

